# PhAGE Enables One-Step Genome Integration of Large DNA Fragments in *Escherichia coli*

**DOI:** 10.64898/2026.04.23.720475

**Authors:** Shingo Nozaki, Yoshihiro Miwa

**Affiliations:** RIKEN BioResource Research Center, 3-1-1 Koyadai, Tsukuba, Ibaraki 305-0074, Japan

## Abstract

*Escherichia coli* is a well-established model organism in molecular biology and biotechnology. Despite its long history as a laboratory workhorse, the efficient single-step chromosomal integration of large DNA fragments remains a challenge. Currently known methods are either simple but have limitations on insert size, or flexible but laborious requiring plasmid construction or multi-step procedures. Here, we present PhAGE (<u>Ph</u>age-<u>A</u>ssisted <u>G</u>enome <u>E</u>ngineering), which enables the integration of ∼20 kb DNA fragments into *E. coli* genome within a single day. PhAGE method uses *in vitro* packaging of recombinant DNA into bacteriophage capsids, followed by general transduction to introduce pre-assembled DNA with flanking homology arms into recipient cells. This approach allows efficient and landing pad-free integration of large constructs into the target loci. We demonstrate its usefulness through rapid integration of multi-gene operons. PhAGE resolves the long-standing trade-off between simplicity and insert size in *E. coli* genome engineering, accelerating strain construction across a wide range of applications, from biosynthetic pathway engineering to genome-scale design.

## Introduction

Genome engineering is an indispensable tool in modern life sciences, supporting progress from basic research to industrial biotechnology. Among microbial systems, *Escherichia coli* serves as a major model organism, providing a diverse platform for genetics, synthetic biology, and metabolic engineering. To support these advances, complex genetic programs are often stably integrated into the genome rather than being maintained on plasmids, ensuring accurate inheritance without continuous antibiotic selection (1, 2). Efficient and precise integration of foreign DNA remains a central requirement for microbial engineering.

In the initial strategies for genome modification in *E. coli*, suicide plasmids were used, which undergo allelic exchange through two successive homologous recombination events (3–5). While this method can incorporate large DNA fragments and generate scarless mutations, it requires multi-step plasmid construction, extended culturing and selection procedures, and extensive screening due to the low frequency of double-crossover. These constraints prompted the development of more efficient genome engineering methods.

A major advance has been made with the λ Red recombinase system (6, 7), which allows simple and efficient one-step gene inactivation using PCR products with short homology arms. While highly effective for small insertions and deletions, its efficiency drops sharply for insertions exceeding ∼3 kb (8), limiting its usefulness for multi-gene constructs. To address this, several strategies have been developed, including homing endonuclease-induced double-strand breaks (8), CRISPR/Cas-assisted integration of large DNA fragments (9–11), and site-specific integration using large serine recombinases (12). While these methods can accommodate larger inserts, they require multi-step workflows or the construction and introduction of multiple plasmids, thereby increasing the complexity of the experiment.

In parallel, general transduction via bacteriophages has long been used to deliver tens of kilobases of genomic DNA between *E. coli* strains (13–15). Phage P1 occasionally package ∼100 kb of host genomic DNA into their capsids, generating transducing particles that deliver these DNA fragments to recipient cells for homologous recombination. However, since the transduced DNA must be derived from the donor E. coli genome, P1 cannot transduce foreign sequences such as gene clusters from other species.

In contrast, λ phage does not mediate general transduction due to its strict *cos*-dependent DNA packaging mechanism (16). Packaging initiates at the *cos* site, restricting λ phage to specialized transduction that transfers only the sequences adjacent to the prophage integration site (17). Despite this limitation *in vivo*, the λ system has been widely used for *in vitro* packaging of recombinant DNA. Pioneering research in the 1970s established that infectious λ particles can be reconstituted *in vitro* from purified heads, tails, and recombinant DNA containing *cos* sites (18, 19). This principle led to the development of λ phage vectors and cosmids capable of cloning DNA fragments up to ∼50 kb (20), which played a central role in early genome mapping and sequencing projects.

Building on these findings, we recently developed iPac (*in vitro* Packaging-assisted DNA assembly), which enables the rapid and accurate construction of large plasmids and phage genomes of up to ∼50 kb (21). A key advantage of iPac is that λ-based packaging inherently enriches constructs of the correct size and effectively eliminates misassembled byproducts. However, iPac is used to introduce episome constructs and phage genomes, not to directly integrate large DNA fragments into the *E. coli* genome.

To overcome these limitations, we developed PhAGE (Phage-Assisted Genome Engineering), a phage-based platform that enables efficient, single-step integration of large DNA fragments into the *E. coli* genome. In this study, we implemented a PhAGE method using λ phage, achieving genome integration within a single day and resolving the long-standing trade-off between simplicity and insert size.

## Materials and Methods

### *E. coli* and phage strain

*E. coli* and λ phage strains used in this study were listed in table S1. SN1171 and SN1187 were obtained from National BioResource project *E. coli*.

### Plasmids

Plasmids used in this study were listed in Table S2. pKD46 and pKD3 (7) were obtained from *E. coli* Genetic Resource Center. pUC19-cos and pUC19-ccdB-cos were constructed by iVEC method with host *E. coli* strain SN1187 (5) using primer sets and templates listed in Tables S3 and S4. For cloning of *pigA-N* operon of *Serratia marcescens*, iPac method were used (21), using primer sets and templates listed in Table S3 and S4. The plasmids constructed in this work were deposited to RIKEN BRC.

### PCR

KOD One DNA polymerase (TOYOBO, Japan) was used for PCR according to the manufacturer’s instructions. PCR products intended for assembly were purified with the NucleoSpin Gel and PCR Clean-up kit (MACHEREY-NAGEL, Germany). Oligonucleotide primers, primer sets and templates used for PCR are listed in Tables S3, S4 and S5. The genomic DNA of *Chromobacterium violaceum* JCM1249T, *Aliivibrio fischeri* JCM18803T and *Serratia marcescens* JCM1239T used as the PCR templates was provided by the RIKEN BRC through the National BioResource Project of the MEXT, Japan (cat. JGD07721, JGD12630 and JGD15966, respectively).

### DNA assembly

DNA assembly using Exo III (XthA) was performed as previously described (21). Combinations of PCR fragments used for DNA assembly were listed in Table S6. Unless otherwise noted, DNA fragments were designed to contain 40-50 bp overlaps at their ends. DNA fragments were adjusted to equal molar concentrations (6 nM each). A 2.5 μL aliquot of the DNA mixture was combined with 2.5 μL of assembly mix (20 mM Tris-HCl (pH 7.9), 100 mM NaCl, 20 mM MgCl2, 10% PEG8000, 2 mM dithiothreitol, and 1.2 U/μL exonuclease III (Takara, Japan)) and gently mixed. The reaction was immediately incubated at 75 °C for 1 min and then left at room temperature for 5 min.

### *In vitro* packaging and transduction

*In vitro* packaging was performed using λ phage packaging extract, Lambda INN (Nippon Gene, Japan). One microliter of the Exo III-assembled DNA mixture was added to 5 μL of packaging extract and incubated at 25 ℃ for 90-120 min. Subsequently, 100 μL of an overnight culture of *E. coli* SN1171, concentrated twofold in 10 mM MgSO₄, was added and incubated for 15 min at room temperature to allow transduction. The entire mixture was plated onto LB agar containing 15 μg/mL chloramphenicol. After incubation at 37 ℃, colonies were counted, and the proportion of correctly assembled clones was determined by colony PCR using primer sets targeting the insertion site listed in Table S5.

### Genome integration by λ Red recombinase

Electrocompetent *E. coli* SN1171 harboring pKD46 was prepared as follows. An overnight culture (1 mL) was inoculated into 100 mL of LB medium supplemented with 100 μg/mL ampicillin and 0.2% arabinose and incubated at 30 °C until the culture reached an OD600 of 0.5. The culture was chilled on ice, and cells were harvested by centrifugation and washed twice with ice-cold 10% glycerol. The cell pellet was resuspended in 1 mL of 10% glycerol and aliquoted into 100 μL portions. Aliquots were frozen in liquid nitrogen and stored at -80 °C until use. For electroporation, a thawed aliquot of competent cells was mixed with 100 ng of linear DNA containing 50 bp homology arms at both ends. Electroporation was performed using an ELEPO21 electroporator (NEPA GENE, Japan) with a 1 mm gap cuvette at 1600 V for 3.5 ms, followed by three additional pulses at 100 V for 50 ms, according to the manufacturer’s instruction. After the pulses, 1 mL of SOC medium was added, and the cells were incubated at 37 °C for 30 min. The culture was then plated onto LB agar containing 15 μg/mL chloramphenicol and incubated at 37 ℃ overnight.

### Bioluminescent colony imaging

Bioluminescent *E. coli* colonies expressing the *lux* operon were imaged in a dark room using a Sony α7CII camera equipped with an FE 28-60 mm F4-5.6 lens. Images were acquired at 60 mm, F5.6, ISO 12800, and a 20-s exposure time.

## Results

### Concept of the PhAGE system

We aimed to develop a method that combines the simplicity of one-step exogenous DNA introduction, as achieved by the λ Red recombinase system, with the capacity to stably integrate large DNA fragments into the *E. coli* genome as in generalized transduction. Building on our previously developed iPac method (21), which enables rapid assembly and bacteriophage-based *in vitro* delivery of large plasmids and phage genomes, we sought to expand its utility to genome engineering. We hypothesized that replacing the standard iPac payload with recombinant construct flanked by long homology arms would mimic the principle of generalized transduction, enabling direct genomic integration of large DNA fragments. This concept underlies the PhAGE system, which couples bacteriophage-based *in vitro* packaging with homologous recombination to achieve efficient, single-step genomic integration (Fig. 1).

**Figure 1.**
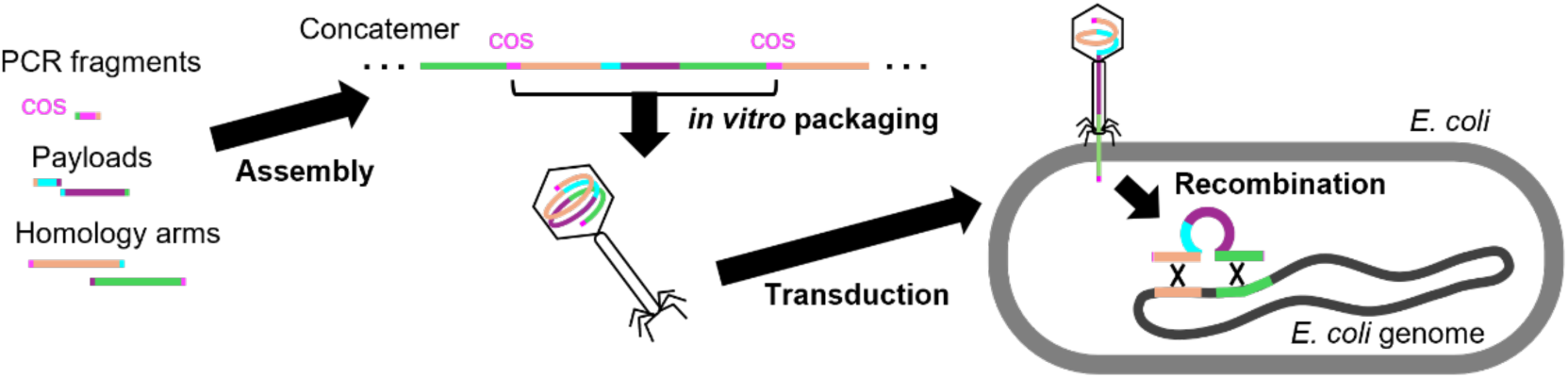
Schematic overview of the PhAGE workflow. PCR fragments, one containing the left homology arm, another containing the right homology arm, several corresponding to the payload sequence, and an additional fragment carrying the λ cos site, were assembled using ∼30-50 bp overlaps. The assembled DNA was packaged *in vitro* into bacteriophage capsids and delivered into *E. coli*. Following delivery, homologous recombination mediated by the flanking homology arms directed integration of the construct into the *E. coli* genome.

### Proof-of-concept of the PhAGE system in *E. coli*

To demonstrate the feasibility of the PhAGE system, we performed a proof-of-concept experiment in *E. coli*. Four DNA payloads (Fig. 2A) were targeted the *yajR* locus in the (Fig. 2B). The smallest payload, *cat* (1 kb), confers chloramphenicol resistance. The other payloads consisted of *cat* positioned upstream of subsets of the *vio* operon from *Chromobacterium violaceum*, which encodes the five enzymes VioABCDE responsible for biosynthesis of violacein, a purple bisindole pigment (22, 23). These constructs formed single operons of increasing size: *cat*-*vioA* (2.6 kb), *cat*-*vioAB* (5.6 kb), and *cat*-*vioABCDE* (hereafter *cat-vio*, 9.2 kb). All payloads were flanked by ∼18 kb homology arms, ensuring that the total construct size exceeded the lower limit of the λ phage packaging window (∼38 kb).

**Figure 2.**
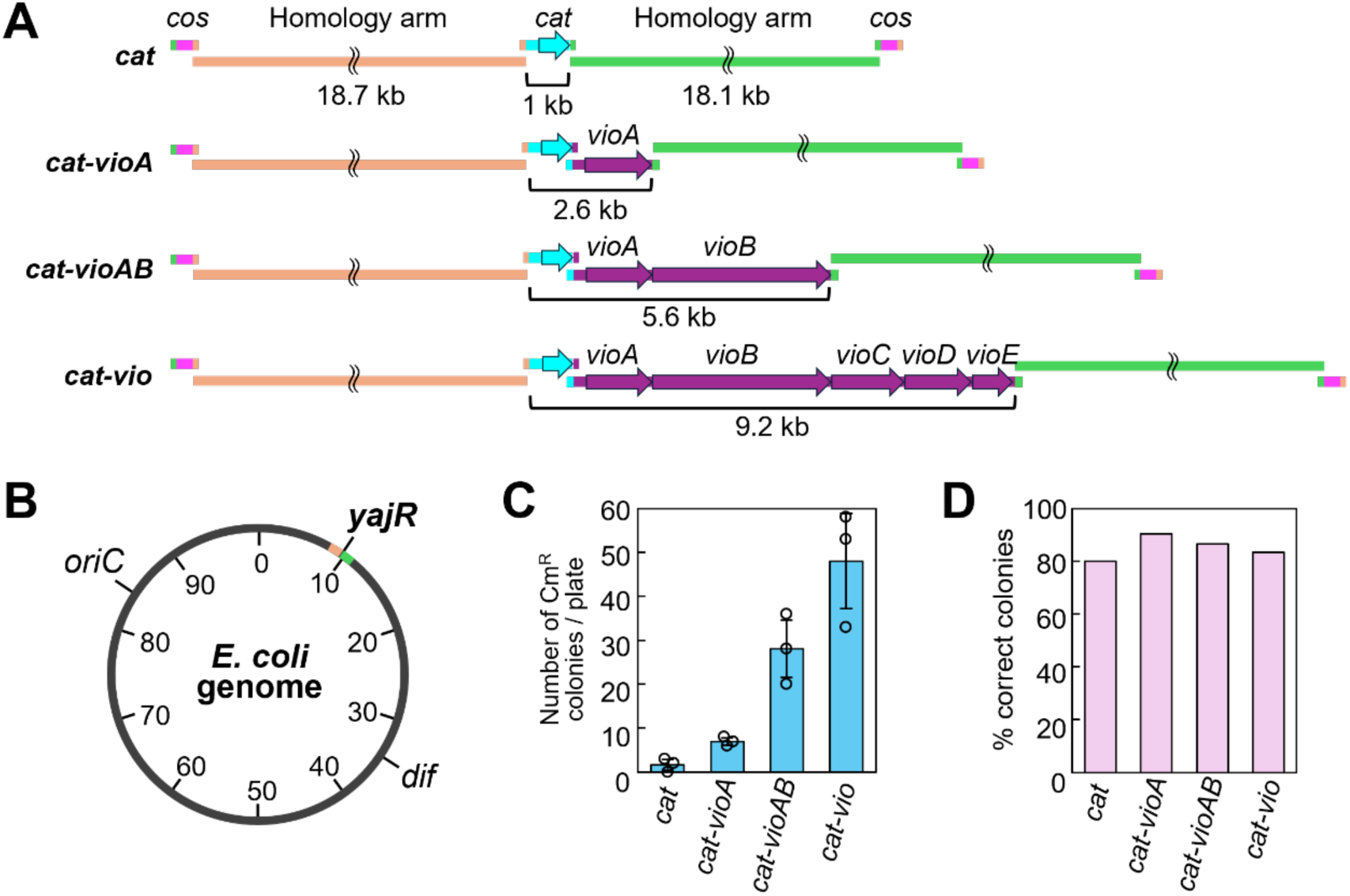
Proof-of-concept of the PhAGE system in *E. coli*. **(A)** Design of the DNA assembly. The payloads comprise *cat* (1 kb), *cat-vioA* (2.6 kb), *cat-vioAB* (5.6 kb), and *cat-vio* (9.2 kb). In each construct, the payload is flanked on both sides by ∼18 kb left and right homology arms corresponding to the *yajR* locus in the *E. coli* genome. A λ phage packaging site (*cos*) is positioned at each end of the construct, which was designed to form concatemers during packaging. **(B)** Location of the *yajR* locus in the *E. coli* genome targeted for integration. **(C)** Average number of chloramphenicol-resistant (Cm^R^) colonies obtained after transduction of each payload into *E. coli*. Error bars indicate the standard deviation, and circles indicate the individual values from biological triplicates. **(D)** Percentage of colonies with correct integration at the target locus, as determined by colony PCR analysis (Fig. S2A-D).

Each DNA fragment was prepared by PCR (Fig. S1). The constructs were assembled using Exonuclease III, packaged into λ phage particles, and delivered into *E. coli*. Transduction yielded an average of 2, 7, 28, and 48 chloramphenicol-resistant (Cm^R^) colonies for *cat* (1 kb), *cat*-*vioA* (2.6 kb), *cat*-*vioAB* (5.6 kb), and *cat*-*vio* (9.2 kb), respectively (Fig. 2C). Because Cm^R^ selection does not guarantee correct insertion, we assessed integration accuracy by colony PCR. This analysis confirmed correct integration at the target genomic locus in 80% (4/5), 90% (19/21), 87% (26/30), and 83% (25/30) of these colonies (Fig. 2D, Fig. S2A-D).

The smallest payload, *cat* (1 kb), yielded few Cm^R^ colonies, likely because its total construct size (38 kb) is close to the lower limit of the λ packaging window (∼38-52 kb). In contrast, larger constructs with total sizes of 39.6 (*cat-vioA*), 42.6 (*cat-vioAB*), and 46.2 kb (*cat-vio*) produced substantially more colonies while maintaining high accuracy.

This trend may reflect more efficient packaging of DNA lengths approaching the native λ genome size (48.5 kb), to which the λ packaging machinery is thought to be evolutionarily optimized.

To benchmark PhAGE against conventional methods, we performed λ Red recombineering using the same payloads with 50 bp homology arms (Fig. S3A, B). This yielded an average of 41, 4, 1, and 2 Cm^R^ colonies for *cat*, *cat*-*vioA*, *cat*-*vioAB*, and *cat*-*vio*, respectively (Fig. S3C). Colony PCR analysis confirmed correct integration in 100% (30/30), 91% (10/11), and 75% (3/4) of the colonies obtained for the three smaller constructs. However, none of the 6 colonies (0/6, 0%) obtained for the 9.2 kb *cat*-*vio* construct carried a correct integration (Fig. S3D).

Together, these results demonstrate that PhAGE enables efficient, single-step integration of DNA fragments from 1 kb to 9.2 kb, outperforming λ Red recombineering for larger payloads. This represents the first demonstration of combining bacteriophage-based *in vitro* packaging with homologous recombination for rapid and precise genome engineering.

### Sequence modification via overlap design

Although complete integration of the *vioABCDE* operon was expected to yield the characteristic purple pigmentation, the *cat-vio* integration strain did not display such distinct pigmentation. We hypothesized that this was due to the operon architecture, in which the upstream *cat* promoter drove expression of *cat* and *vioABCDE* (Fig. 3A), resulting in insufficient expression of the *vio* operon.

**Figure 3.**
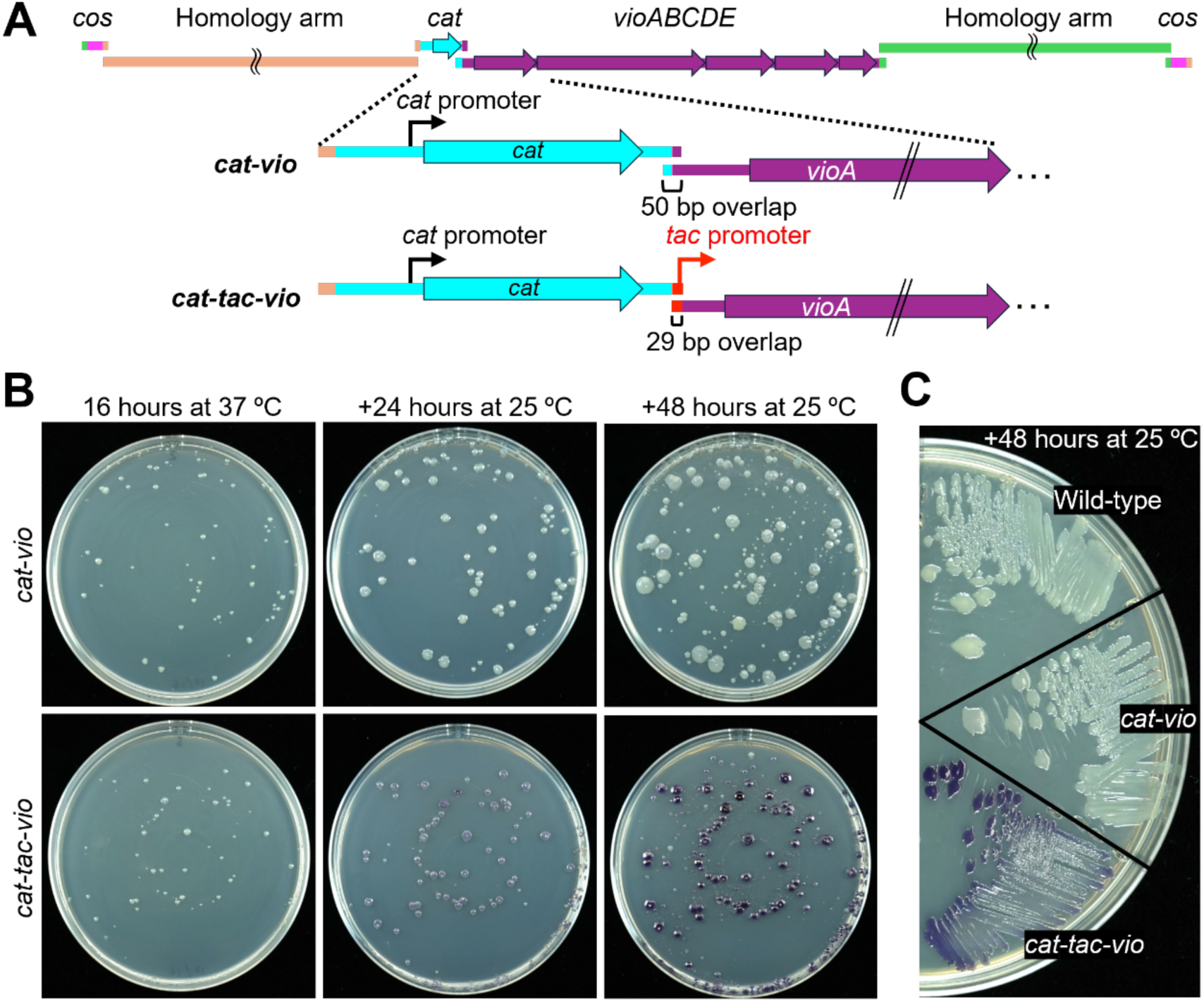
*tac* promoter insertion via overlap design. **(A)** Schematic illustration of the constructs, defined as *cat-vio* (without an additional *tac* promoter) or *cat-tac-vio* (with an additional *tac* promoter inserted within the terminal overlap between the *cat* and *vioABCDE* fragments). **(B)** Plate images of *cat-vio* and *cat-tac-vio* integration strains obtained via PhAGE, showing differences in colony pigmentation after incubation at the indicated time and temperature. **(C)** Comparison of colony color for the wild-type, *cat-vio*, and *cat-tac-vio* strains grown on the same plate to allow direct visual comparison.

To test this hypothesis, and to demonstrate that overlap design enables precise sequence modification at the fragment junction, we inserted an additional *tac* promoter (24) into the terminal overlap between the *cat* and *vioABCDE* fragments (Fig. 3A). We then examined whether this modification enhanced expression of the *vio* operon.

After introducing these DNA constructs via PhAGE and incubating the plates containing chloramphenicol at 37 °C for 16 hours, colonies of the *cat-tac-vio* integration strain exhibited faint purple pigmentation (Fig. 3B). When the plates were further incubated at 25 °C for an additional 24 hours, the purple pigmentation became clearly visible, and after 48 hours, the colonies developed a markedly deeper purple (Fig. 3B, C). In contrast, the *cat-vio* strain lacking the additional promoter exhibited a barely perceptible pigmentation, comparable to the wild-type strain (Fig. 3C).

These results demonstrate that PhAGE enables seamless assembly and integration of DNA fragments, and that small insertions or deletions can be readily introduced by appropriately designing the overlap regions.

### Targeted integration at various genomic loci

To evaluate the versatility of PhAGE for precise genome engineering at different chromosomal positions, we selected four loci in the *E. coli* genome, *yieL*, *yajR*, *ypjA*, and *yneO*, located near the origin of replication (*oriC*), in the right chromosomal arm (the same site targeted in previous experiments), in the left chromosomal arm, and near the *dif* site, respectively (Fig. 4A).

**Figure 4.**
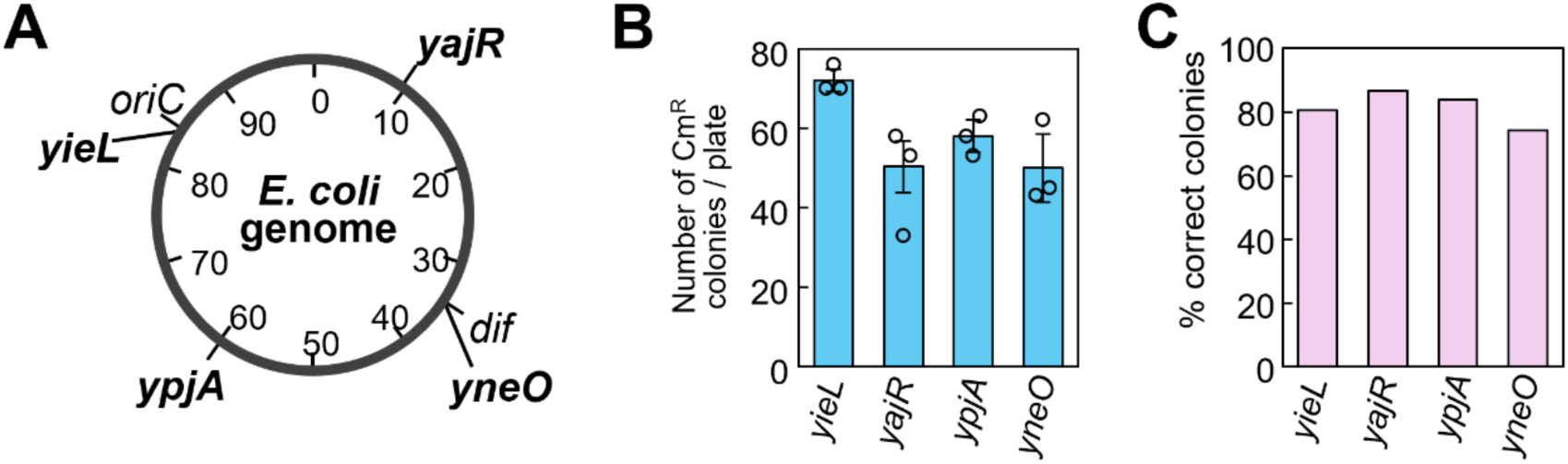
Targeted integration at various genomic loci. **(A)** Genomic map indicating the four target loci for integration: *yieL* (near *oriC*), *yajR* (right chromosomal arm), *ypjA* (left chromosomal arm), and *yneO* (near the *dif* site). **(B)** Average number of Cm^R^ colonies obtained following PhAGE-mediated integration at each target locus. Error bars indicate the standard deviation, and circles indicate individual values from biological triplicates. **(C)** Proportion of correct integration at each locus, determined by colony PCR analysis of randomly selected 30 or 31 colonies obtained after PhAGE-mediated integration.

Introducing the *cat-tac-vio* construct into these sites yielded average numbers of Cm^R^ colonies of 72, 50, 58, and 50, respectively (Fig. 4B). The slightly higher colony number observed at *yieL* may reflect the increased copy number of *oriC*-proximal regions during active replication. Colony PCR analysis confirmed correct integration rates of 81% (25/31), 87% (26/30), 84% (26/31), and 74% (23/31) for integrations at *yieL*, *yajR*, *ypjA*, and *yneO*, respectively (Fig. 4C).

Overall, integration efficiencies remained high across all tested loci, indicating that chromosomal position had only a minor effect on PhAGE performance. These results demonstrate that PhAGE enables efficient, targeted integration across diverse genomic loci, and that altering the PCR-amplified homology arms is sufficient to retarget integration into new chromosomal positions.

### Analysis of incorrect integration events

To better understand the limitations of PhAGE-mediated integration, we examined colonies in which integration was unsuccessful. PCR analysis of these colonies revealed not only the wild-type band but also a faint band corresponding to the intended insert (Fig. S2A-D; see also Fig. 5B). Because the donor DNA enters *E. coli* with complementary single-stranded *cos* termini, we hypothesized that these ends could anneal to form circular molecules, which might subsequently integrate into the chromosome in an unintended manner (Fig. 5A).

**Figure 5.**
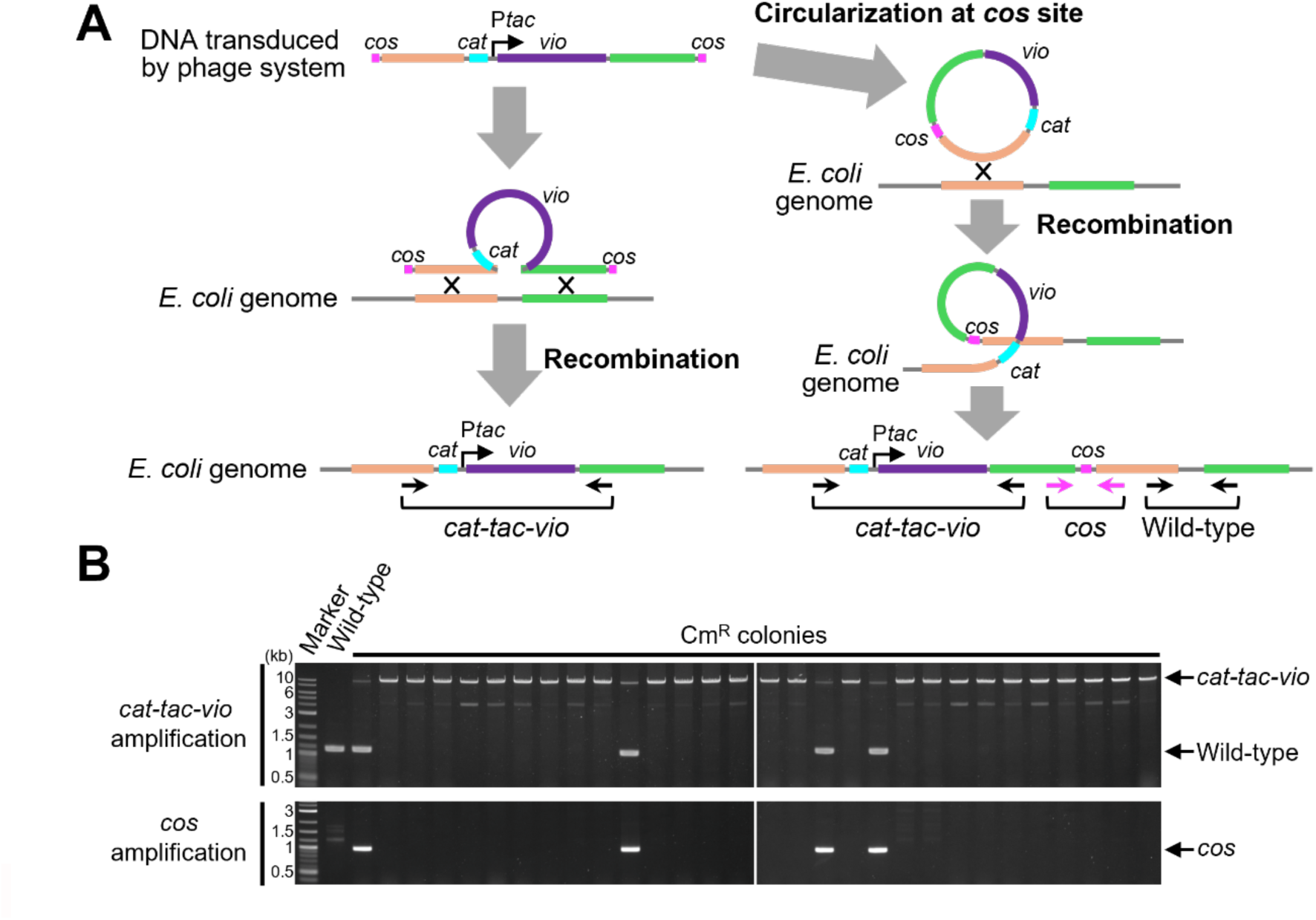
Mechanism of incorrect integration. **(A)** Schematic representation of the integration pathways. The left pathway illustrates the correct integration event, whereas the right pathway depicts an incorrect event in which donor DNA undergoes circularization at the *cos* sites prior to integration, resulting in a merodiploid state. Black arrows indicate primer sets used to detect either the *cat-tac-vio* cassette or the wild-type locus, while magenta arrows indicate primer sets specifically designed to detect circularization at the *cos* sites. **(B)** Agarose gel electrophoresis of colony PCR products. Colony PCR using the two primer sets described in Fig. 5A was performed on 30 randomly selected Cm^R^ colonies obtained after PhAGE-mediated integration. Wild-type *E. coli* was included as a negative control. The upper panel shows PCR results obtained with the *cat-tac-vio* primer set, whereas the lower panel shows PCR results obtained with the primer set specific for *cos*-site circularization. Arrows indicate the bands corresponding to the *cat-tac-vio* cassette, the wild-type allele, and the *cos*-site circularization product, as labeled.

To test this possibility, we performed colony PCR using a primer set designed to yield amplification only when circularization at the *cos* sites had occurred (Fig. 5A). Among the Cm^R^ colonies obtained after PhAGE-mediated integration, the circularization-specific PCR band was detected exclusively in colonies lacking correct integration, and all such colonies produced the band (Fig. 5B). This confirms that circularization of donor DNA at the *cos* sites underlies the incorrect integration events observed.

### Design and validation of a failure-prevention strategy

Despite the competing circularization events, PhAGE already achieves a high success rate, with approximately 80% of colonies containing the desired insertion. Thus, the method reliably yields correct integrants, although further increasing the proportion of desired colonies would enhance its robustness.

To address this, we designed a strategy to selectively eliminate cells that carry circularized donor DNA. Specifically, the *ccdB* toxin gene, previously used as a counter-selection marker (25), was placed adjacent to the *cos* site at the termini of the donor DNA. When circularization does not occur, the linear DNA ends are expected to be degraded by endogenous exonucleases such as RecBCD (26). This degradation removes the *ccdB* as well as *cos,* allowing the host cell to survive and ensuring correct integration of the payload. In contrast, when circularization occurs, the *cos* ends anneal and protect *ccdB* from degradation, resulting in retention of the toxin gene and subsequent cell death (Fig. 6A). This design was therefore expected to eliminate colonies derived from circularized integration events and enrich for correct integrants.

**Figure 6.**
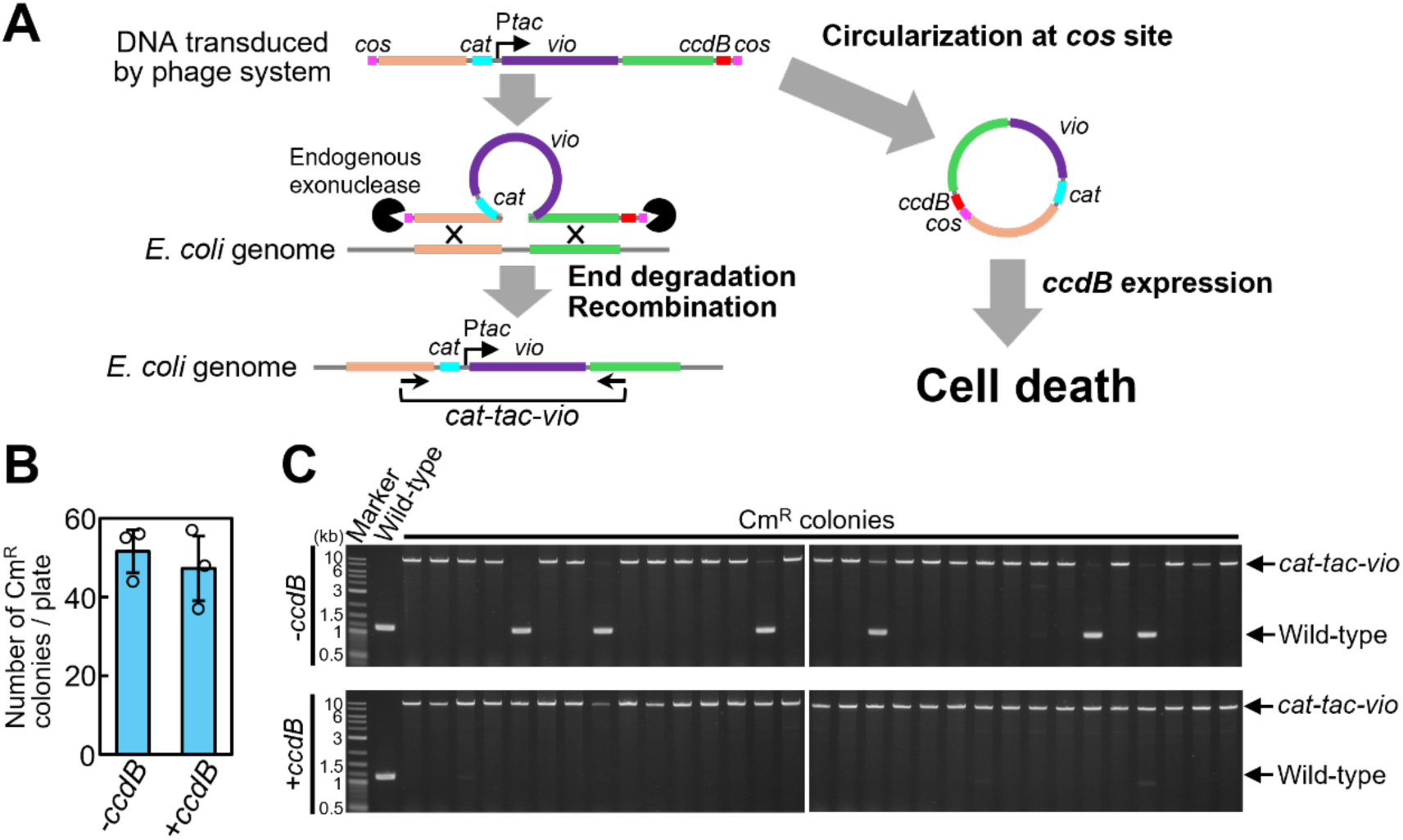
Design and validation of a failure-prevention strategy. **(A)** Schematic design of the failure-prevention strategy using *ccdB*. The *ccdB* gene was positioned adjacent to the *cos* site of the donor DNA. Correct integration leads to degradation of terminal sequences and removal of both the *cos* site and *ccdB*, allowing host survival (left), whereas incorrect circularization preserves both cos sites and *ccdB*, resulting in toxicity and cell death (right). **(B)** Comparison of colony numbers with and without *ccdB*. Average Cm^R^ colony counts following PhAGE-mediated integration with or without *ccdB*. Error bars indicate standard deviation, and circles represent individual values from biological triplicates. **(C)** Agarose gel electrophoresis of colony PCR products. Colony PCR was performed on 31 randomly selected Cm^R^ colonies following PhAGE-mediated integration, for both constructs without and with the *ccdB* gene, using the *cat-tac-vio* primer set. Wild-type *E. coli* served as a negative control. The upper panel shows PCR results without *ccdB*, while the lower panel shows results with *ccdB*. Arrows indicate the bands corresponding to the *cat-tac-vio* cassette and the wild-type allele.

To validate this concept, we compared PhAGE with and without *ccdB*. Inclusion of *ccdB* caused a slight decrease in the average colony number, from 52 to 47 (p = 0.57 by Welch’s t-test) (Fig. 6B). Importantly, the proportion of desired colonies increased from 80.6% (25/31) to 100% (31/31) (Fig. 6C). These results demonstrate that the failure-prevention design effectively eliminates incorrect integration events and enriches for correctly integrated colonies.

### Application to other payloads

To further evaluate the versatility of PhAGE, we replaced the *vio* gene cluster with alternative genetic payloads. As a first example, the *luxCDABEG* operon from *Aliivibrio fischeri*, which encodes luciferase and the enzymes required for luciferin biosynthesis (27), was introduced into the *yajR* locus. This resulted in a 7.7 kb *cat*-*tac*-*lux* cassette (Fig. 7A). Colonies obtained after PhAGE-mediated integration exhibited visible bioluminescence in the dark, consistent with luciferin-luciferase reaction activity (Fig. 7B). Colony PCR confirmed that correct integration in 73% (11/15) of colonies (fig. 7C). Next, a substantially larger payload, the *pigA-N* operon from *Serratia marcescens*, which encodes the enzymes responsible for biosynthesis of the red pigment, prodigiosin (28), was similarly integrated into the *yajR* locus. The resulting cat-tac-pig cassette was 22.1 kb in size, so the homology arms were adjusted to 11.3 kb and 11.5 kb to keep the total construct within the λ phage packaging limit. Because the *pigA-N* operon itself spans 21.2 kb, the PCR product was divided into two fragments of 11.3 kb and 9.9 kb to facilitate reliable amplification (fig.7D). To prevent incorrect integration events, we also tested a circularization-blocking strategy using *ccdB*.

**Figure 7.**
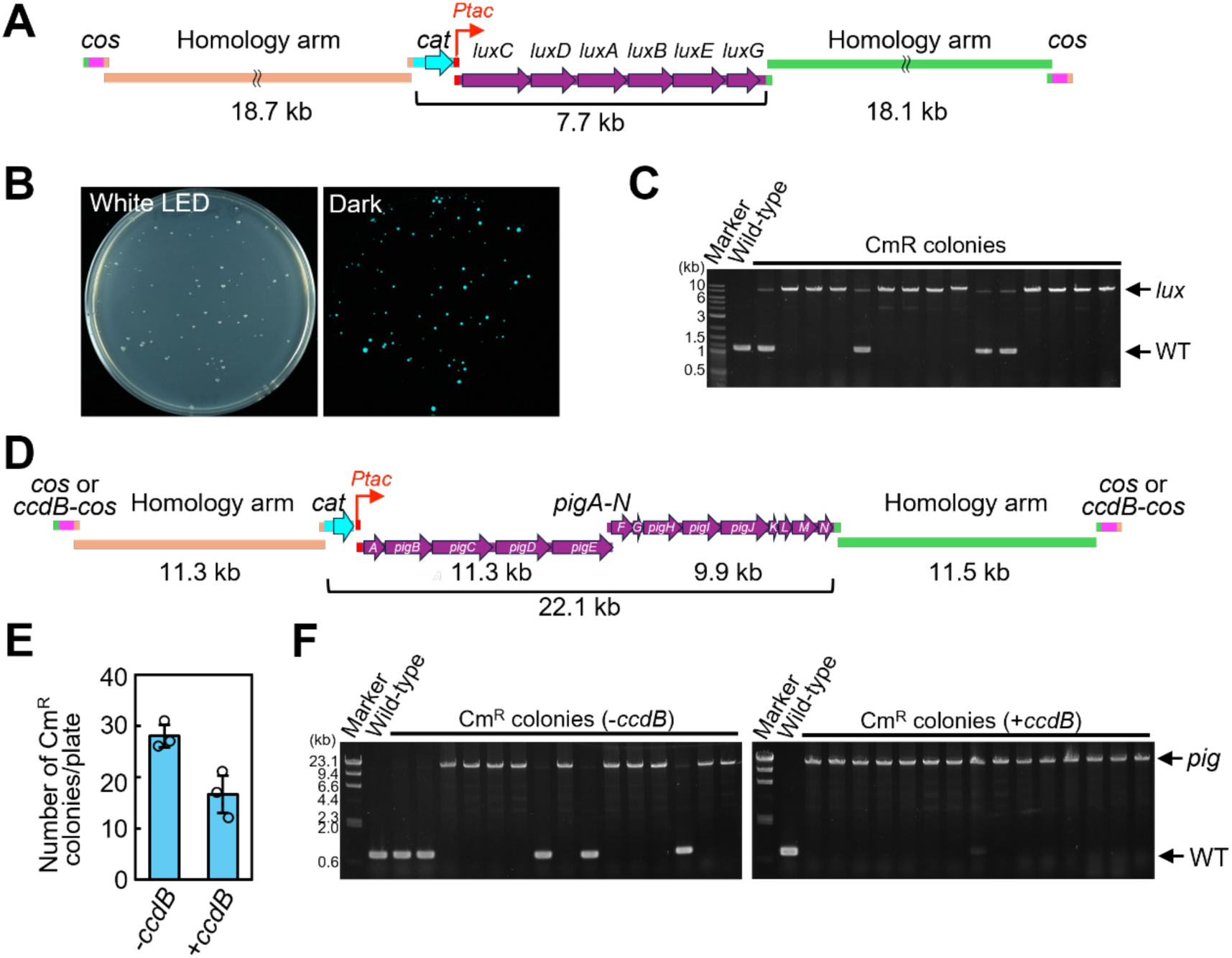
PhAGE-mediated integration of diverse genetic payloads. **(A)** Schematic representation of PCR products used for integration of the *luxCDABEG* operon from *Aliivibrio fischeri* into the *yajR* locus. **(B)** Representative plate images after PhAGE-mediated integration of the *lux* operon. Colonies were photographed under white LED illumination (left) and in the dark (right), showing bioluminescent colonies. **(C)** Colony PCR analysis of *lux* integration. Wild-type *E. coli* served as the negative control. Arrows indicate the bands corresponding to the *lux* cassette and the wild-type allele. **(D)** Schematic representation of PCR products used for integration of the *pigA-N* operon from *Serratia marcescens* into the *yajR* locus, using either cos or *ccdB*-*cos* termini. **(E)** Average number of Cm^R^ colonies obtained after PhAGE-mediated integration of the *pig* operon. Error bars indicate standard deviation, and circles represent individual values from biological triplicates. **(F)** Colony PCR analysis of *pig* integration. Results are shown for conditions without *ccdB* (left) and with *ccdB* (right). Wild-type *E. coli* served as the negative control. Arrows indicate the bands corresponding to the *pig* cassette and the wild-type allele.

PhAGE-mediated integration yielded an average of 28 Cm^R^ colonies without *ccdB* and 17 colonies with *ccdB* (p=0.029, Welch’s t-test) (Fig. 7E). Colony PCR analysis revealed that the proportion of correctly integrated colonies was 67% (10/15) without *ccdB*, whereas all colonies (15/15; 100%) were correctly integrated when *ccdB* was included (Fig. 7F). These results demonstrate that PhAGE can support one-step chromosomal integration of DNA cassettes as large as 22.1 kb.

Colonies harboring the genome-integrated *pigA-N* operon did not exhibit red pigmentation under the tested conditions, whereas expression from a multicopy pUC19-based plasmid carrying the same operon (pUCcos-pig) displayed the expected pigmentation (Fig. S5). This suggests that the lack of pigmentation in genome-integrated strains is due to insufficient expression from the single-copy locus, rather than defects in the integrated construct.

Together, these results highlight the ability of PhAGE to achieve one-step targeted integration of diverse genetic payloads, including fragments up to 22.1 kb, into *E. coli* genome.

## Discussion

### Conceptual and Practical Advances of the PhAGE System

PhAGE extends our previously established iPac system from circular DNA assembly to chromosomal integration. While iPac assembles circular DNA such as phage genomes or plasmids, PhAGE enables the direct integration of exogenous DNA into the host genome. PhAGE utilizes the unique ability of phages to package and inject large DNA molecules, enabling the efficient, one-step integration of fragments up to 22 kb into the *E. coli* genome. This feature provides a versatile platform for rapidly introducing multiple gene clusters, offering broad potential in synthetic biology and industrial biotechnology.

Many biosynthetic pathways form large, multi-gene modules that play a central role in metabolic engineering applications (29). PhAGE provides a practical method for directly introducing these modules into the genome by enabling the integration of large constructs in a single step.

Another advantage of the PhAGE is that it requires minimal preparation on the host side. In contrast to electroporation or methods that require the prior plasmid introduction, λ phage transduction can be performed using simple overnight cultures treated with Mg^2+^. This eliminates the need to prepare special competent cells, lowering the barrier to routine genome engineering.

### Potential Payload Limits and Future Extensions of the System

In this study, we achieved successful integration of a 22 kb payload using ∼10 kb homology arms. Classical RecA-dependent recombination studies indicate that homology arms of at least 1 kb substantially enhance recombination efficiency (30), suggesting arms of several kilobases, such as ∼5 kb, are likely sufficient. Although the exact upper limit has not yet been determined, given that the λ packaging capacity is ∼52 kb, a payload of around 30-40 kb may be the practical maximum for this system. Importantly, this limit is imposed by the λ packaging system rather than by the PhAGE strategy itself. Alternative phage systems with larger packaging capacities or different host ranges could in principle extend the payload size and the host range spectrum, enabling broader applications.

### Modular Use of *cos*-Proximal Regions for Transient Factor Delivery in PhAGE

PhAGE-mediated integration relies on long homology arms and degradation of linear DNA ends by host exonucleases. As a result, sequences placed adjacent to the *cos* sites are eliminated during recombination. We exploited this property by positioning *ccdB* in the *cos*-proximal region to prevent undesirable circular DNA integration. More broadly, these regions may serve as modular sites for incorporating transiently acting factors such as recombinases, CRISPR-based modules, or recombination enhancers. Embedding such factors directly within the disposable portion of the packaged DNA could, in principle, eliminate the need for host strains pre-equipped with accessory plasmids, simplifying strain preparation and expanding the applicability of the system. PhAGE therefore offers a potential route for delivering both transient functions, in a manner analogous to phage early genes that rapidly reprogram the host environment immediately after infection, and permanently integrated genetic elements in a single step.

### Potential Extensions of the PhAGE System

Recent studies have shown that engineered bacteriophages can mediate *in situ* genome editing of gut-resident bacteria (31), highlighting the feasibility of phage-based genetic manipulation directly within complex microbial communities. Given the minimal host preparation required and the conceptual ability to co-deliver transient editing factors, it is tempting to speculate that the PhAGE framework could be adapted for *in situ* genome rewriting of gut microbiota. Achieving this will require addressing several practical challenges, including the limited yield of phage particles produced by *in vitro* packaging. Incorporating *in vivo* packaging strategies, such as high-efficiency cosmid packaging in bacterial hosts (32, 33), could substantially increase particle yield and improve feasibility for microbiome applications.

Together, these considerations suggest that PhAGE, whether implemented through *in vitro* or *in vivo* packaging, offers a flexible foundation for deploying programmable genome engineering tools within complex microbial environments.

### Concluding Remarks on PhAGE

In summary, PhAGE is a practical solution for rapid integration of large DNA into *E. coli* genome, overcoming the long-standing trade-off between simplicity and insert size. These advances provide a solid foundation for future methodological development and expand the toolkit available for bacterial genome engineering.

## Acknowledgement

We gratefully acknowledge the NBRP *E. coli* (Japan) and the *E. coli* Genetic Resource Center (CT, USA) for providing *E. coli* strains, and the RIKEN DNA Bank (Japan) for supplying genomic DNA. We also thank Dr. Tetsushi Iida for DNA sequencing and Miharu Hata for technical assistance with PCR and gel electrophoresis. Finally, we appreciate the members of our laboratory for maintaining a pleasant working environment.

## Author contributions

S.N. conceived the project. Y.M. supervised the project. S.N. designed and conducted experiments. S.N. wrote the manuscript.

## Reference

1. Zucca, S., Pasotti, L., Politi, N., Cusella De Angelis, M.G. and Magni, P. (2013) A standard vector for the chromosomal integration and characterization of BioBrick^TM^ parts in *Escherichia coli*. J. Biol. Eng., 7, 12.

2. Yeom, S.-J., Lee, D.-H., Kim, Y.J., Lee, J., Kwon, K.K., Han, G.H., Kim, H., Kim, H.-S. and Lee, S.-G. (2016) Long-term stable and tightly controlled expression of recombinant proteins in antibiotics-free conditions. PloS One, 11, e0166890.

3. Ried, J.L. and Collmer, A. (1987) An nptI-sacB-sacR cartridge for constructing directed, unmarked mutations in gram-negative bacteria by marker exchange-eviction mutagenesis. Gene, 57, 239–246.

4. Hamilton, C.M., Aldea, M., Washburn, B.K., Babitzke, P. and Kushner, S.R. (1989) New method for generating deletions and gene replacements in Escherichia coli. J. Bacteriol., 10.1128/jb.171.9.4617-4622.1989.

5. Nozaki, S. and Niki, H. (2019) Exonuclease III (XthA) enforces *in vivo* DNA cloning of *Escherichia coli* to create cohesive ends. J. Bacteriol., 201, e00660–18.

6. Murphy, K.C. (1998) Use of bacteriophage λ recombination functions to promote gene replacement in *Escherichia coli*. J. Bacteriol., 180, 2063–2071.

7. Datsenko, K.A. and Wanner, B.L. (2000) One-step inactivation of chromosomal genes in *Escherichia coli* K-12 using PCR products. Proc. Natl. Acad. Sci., 97, 6640–6645.

8. Kuhlman, T.E. and Cox, E.C. (2010) Site-specific chromosomal integration of large synthetic constructs. Nucleic Acids Res., 38, e92.

9. Chung, M.-E., Yeh, I.-H., Sung, L.-Y., Wu, M.-Y., Chao, Y.-P., Ng, I.-S. and Hu, Y.-C. (2017) Enhanced integration of large DNA into *E. coli* chromosome by CRISPR/Cas9. Biotechnol. Bioeng., 114, 172–183.

10. Li, Y., Yan, F., Wu, H., Li, G., Han, Y., Ma, Q., Fan, X., Zhang, C., Xu, Q., Xie, X., et al. (2019) Multiple-step chromosomal integration of divided segments from a large DNA fragment via CRISPR/Cas9 in *Escherichia coli*. J. Ind. Microbiol. Biotechnol., 46, 81–90.

11. Fredens, J., Wang, K., de la Torre, D., Funke, L.F.H., Robertson, W.E., Christova, Y., Chia, T., Schmied, W.H., Dunkelmann, D.L., Beránek, V., et al. (2019) Total synthesis of *Escherichia coli* with a recoded genome. Nature, 569, 514–518.

12. Riley, L.A., Payne, I.C., Tumen-Velasquez, M. and Guss, A.M. (2023) Simple and rapid site-specific integration of multiple heterologous DNAs into the *Escherichia coli* chromosome. J. Bacteriol., 205, e0033822.

13. Zinder, N.D. and Lederberg, J. (1952) Genetic exchange in *Salmonella*. J. Bacteriol., 64, 679–699.

14. Lennox, E.S. (1955) Transduction of linked genetic characters of the host by bacteriophage P1. Virology, 1, 190–206.

15. Thomason, L.C., Costantino, N. and Court, D.L. (2007) E. coli Genome Manipulation by P1 Transduction. Curr. Protoc. Mol. Biol., 79, 1.17.1-1.17.8.

16. Hohn, B. (1983) DNA sequences necessary for packaging of bacteriophage lambda DNA. Proc. Natl. Acad. Sci. U. S. A., 80, 7456–7460.

17. Fukumaki, Y., Shimada, K. and Takagi, Y. (1976) Specialized transduction of colicin E1 DNA in *Escherichia coli* K-12. Proc. Natl. Acad. Sci. U. S. A., 73, 3238–3242.

18. Hohn, B., Wurtz, M., Klein, B., Lustig, A. and Hohn, T. (1974) Phage lambda DNA packaging, *in vitro*. J. Supramol. Struct., 2, 302–317.

19. Hohn, B. and Murray, K. (1977) Packaging recombinant DNA molecules into bacteriophage particles *in vitro*. Proc. Natl. Acad. Sci., 74, 3259–3263.

20. Collins, J. and Hohn, B. (1978) Cosmids: a type of plasmid gene-cloning vector that is packageable *in vitro* in bacteriophage lambda heads. Proc. Natl. Acad. Sci., 75, 4242–4246.

21. Nozaki, S. (2022) Rapid and accurate assembly of large DNA assisted by *in vitro* packaging of bacteriophage. ACS Synth. Biol., 11, 4113–4122.

22. August, P.R., Grossman, T.H., Minor, C., Draper, M.P., MacNeil, I.A., Pemberton, J.M., Call, K.M., Holt, D. and Osburne, M.S. (2000) Sequence analysis and functional characterization of the violacein biosynthetic pathway from *Chromobacterium violaceum*. J. Mol. Microbiol. Biotechnol., 2, 513–519.

23. Balibar, C.J. and Walsh, C.T. (2006) *In vitro* biosynthesis of violacein from l-tryptophan by the enzymes VioA−E from *Chromobacterium violaceum*. Biochemistry, 45, 15444–15457.

24. de Boer, H.A., Comstock, L.J. and Vasser, M. (1983) The *tac* promoter: a functional hybrid derived from the *trp* and *lac* promoters. Proc. Natl. Acad. Sci., 80, 21–25.

25. Bernard, P., Gabarit, P., Bahassi, E.M. and Couturier, M. (1994) Positive-selection vectors using the F plasmid *ccdB* killer gene. Gene, 148, 71–74.

26. El Karoui, M., Dabert, P., Gruss, A. and Amundsen, S.K. (1999) Gene replacement with linear DNA in electroporated wild-type *Escherichia coli*. Nucleic Acids Res., 27, 1296–1299.

27. Meighen, E.A. (1991) Molecular biology of bacterial bioluminescence. Microbiol. Rev., 10.1128/mr.55.1.123-142.1991.

28. Harris, A.K.P., Williamson, N.R., Slater, H., Cox, A., Abbasi, S., Foulds, I., Simonsen, H.T., Leeper, F.J. and Salmond, G.P.C. (2004) The *Serratia* gene cluster encoding biosynthesis of the red antibiotic, prodigiosin, shows species- and strain-dependent genome context variation. Microbiology, 150, 3547–3560.

29. Nielsen, J. and Keasling, J.D. (2016) Engineering Cellular Metabolism. Cell, 164, 1185–1197.

30. King, S.R. and Richardson, J.P. (1986) Role of homology and pathway specificity for recombination between plasmids and bacteriophage λ. Mol. Gen. Genet. MGG, 204, 141–147.

31. Brödel, A.K., Charpenay, L.H., Galtier, M., Fuche, F.J., Terrasse, R., Poquet, C., Havránek, J., Pignotti, S., Krawczyk, A., Arraou, M., et al. (2024) *In situ* targeted base editing of bacteria in the mouse gut. Nature, 632, 877–884.

32. Cronan, J.E. (2003) Cosmid-based system for transient expression and absolute off- to-on transcriptional control of *Escherichia coli* genes. J. Bacteriol., 10.1128/jb.185.22.6522-6529.2003.

33. Cronan, J.E. (2023) Two neglected but valuable genetic tools for *Escherichia coli* and other bacteria: *In vivo* cosmid packaging and inducible plasmid replication. Mol. Microbiol., 120, 783–790.

