## Supporting information for "PhAGE Enables One-Step Genome Integration of Large DNA Fragments in *Escherichia coli*"

**Table S1: Bacterial and phage strains**

| **Strain** | **Genotype** | **Reference or Source** |
|---|---|---|
| SN1171 | F-, *rph-1*, Δ*hsdR*, Δ*endA* | Nozaki and Niki, 2019 |
| SN1187 | F-, *rph-1*, Δ*hsdR*, Δ*endA*, Δ*recA* | Nozaki and Niki, 2019 |
| XL-10 Gold | TetR Δ(*mcrA*)183 Δ(*mcrCB-hsdSMR-mrr*)173 *endA1 supE44 thi-1 recA1 gyrA96 relA1 lac Hte* [F'*proAB lacIq*ZΔM15 Tn10 (TetR) *Amy* CamR] | Agilent |
| λ *cI857* | λ phage *cI857* | Nozaki, 2022 |

**Table S2: Plasmids**

| **Name** | **Description** | **Reference** |
|---|---|---|
| pKD3 | *cat* gene cassette | Datsenko and Wanner, 2000 |

| | | |
|---|---|---|
| pKD46 | λ Red recombinase expression | Datsenko and Wanner, 2000 |
| pUC19-cos | Cloned *cos* sequence | This work |
| pUC19-ccdB-cos | Cloned *ccdB-cos* sequence | This work |
| pUCcos-pig | Cloned prodigiosin biosynthesis genes | This work |

**Table S3: Oligonucleotide primers for PCR**

| **Name** | **Oligonucleotide sequence (5' to 3')** |
|---|---|
| ONR1280 | CCTTTTCCATTTCTGAACATTGCCACCACGCACGTTGTGATATG |
| ONR1281 | CGCCCATCAGGTAACGCGGTTTGTCCATTGTTCATTCCACGGACA |
| ONR1278 | TCGCCGGGATTTCAATGCGCAAACTGCTGACATACGGCGGTTCTTTCATGC ATATGAATATCCTCCTTAG |
| ONR1099 | TTAGTTCAACCGGACTCATTCATCAGTGTAGGCTGGAGCTGCTTC |
| ONR1102 | ACTTCGAAGCAGCTCCAGCCTACACTGATGAATGAGTCCGGTTGA |
| ONR1105 | GTCTTTGTCTTCTGCGTCGGGTTGATCGTGGTTGCGGAAATTGTGTTGTGTG TGTTTCCCGAACGGTCAT |
| ONR1104 | GTCTTTGTCTTCTGCGTCGGGTTGATCGTGGTTGCGGAAATTGTGTTGTGTC AGGCCTCTCTAGAAAGCT |
| ONR1103 | GTCTTTGTCTTCTGCGTCGGGTTGATCGTGGTTGCGGAAATTGTGTTGTGGA CAACTGAGCGGCTACTCG |
| ONR1163 | GTCTTTGTCTTCTGCGTCGGGTTGATCGTGGTTGCGGAAATTGTGTTGTGGT GTAGGCTGGAGCTGCTTC |
| ONR1282 | GTTTTTGTCCGTGGAATGAACAATGGACAAACCGCGTTACCTGAT |
| ONR1097 | CATGAAAGAACCGCCGTATG |

| | |
|---|---|
| ONR1100 | CACAACACAATTTCCGCAAC |
| ONR1283 | CATCTACATATCACAACGTGCGTGGTGGCAATGTTCAGAAATGGA |
| ONR1284 | CATTATACGAGCCGATGATTAATTGTCAAGTGTAGGCTGGAGCTGCTTC |
| ONR1272 | TTGACAATTAATCATCGGCTCGTATAATGCATGACGAGGAAACGAAAGGCAG |
| ONR1279 | GTCTTTGTCTTCTGCGTCGGGTTGATCGTGGTTGCGGAAATTGTGTTGTGGACAACTGAGCGGCTACTCG |
| ONR1262 | CTACACCCGCACCACCGAACAGACCAACTTTACCGCCCTTCCACGCACGTTGTGATATGT |
| ONR1263 | ATGGACACCGAAGCCATGGGTGATTAAAGAGGCCGGATTGCATTGTTCATTCCACGGACA |
| ONR1268 | ATATTCACCCGATGACCAAAGACGAAGCAGGAGTCTGGTCCATATGAATATCCTCCTTAG |
| ONR1269 | TTTTCCCTTCAGCAAGCAGGTTATCCATGATTTGCGGGATGACAACTGAGCGGCTACTCG |
| ONR1264 | CAATCCGGCCTCTTTAATCA |
| ONR1265 | ATCCCGCAAATCATGGATAA |
| ONR1266 | GACCAGACTCCTGCTTCGTC |
| ONR1267 | AAGGGCGGTAAAGTTGGTCT |
| ONR1201 | CGGCATCCGGTATACCGGTTTTGTTCAGATGCTCCTGCACCCACGCACGTTGTGATATGT |
| ONR1202 | TACAAGCACATCGCGGATAACGTCTGTCAATGGTTGGATGCATTGTTCATTCCACGGACA |

| | |
|---|---|
| ONR1203 | AAGATGTTGGCACTGTACTCAATAGTGCTGGCACCCAAACCATATGAATATCCTCCTTAG |
| ONR1204 | TTGCCTCTCCGTTAGCTGAAACTGTTTGTATTCCGCCGTCGACAACTGAGCGGCTACTCG |
| ONR1197 | CATCCAACCATTGACAGACG |
| ONR1198 | GACGGCGGAATACAAACAGT |
| ONR1199 | GTTTGGGTGCCAGCACTATT |
| ONR1200 | GTGCAGGAGCATCTGAACAA |
| ONR1191 | GATAGAACGCAATCAATGCCGCTAATGCGAAAGTAAGGCGCCACGCACGTTGTGATATGT |
| ONR1192 | TGAAGAGATGACGGTCTATGCTCCTGTCCCTGTACCCGTACATTGTTCATTCCACGGACA |
| ONR1193 | ACGGTCAACGGTATTGAACTCATTGAGGTTGGCGGTAATTCATATGAATATCCTCCTTAG |
| ONR1194 | CCGGATCCACAACAGGGGGATTATTGATGGGATCGGGTGTGACAACTGAGCGGCTACTCG |
| ONR1187 | TACGGGTACAGGGACAGGAG |
| ONR1188 | ACACCCGATCCCATCAATAA |
| ONR1189 | AATTACCGCCAACCTCAATG |
| ONR1190 | CGCCTTACTTTCGCATTAGC |
| ONR1173 | TTGACAATTAATCATCGGCTCGTATAATGCGGTTGTATTGTCTATGCCT |
| ONR1174 | GTCTTTGTCTTCTGCGTCGGGTTGATCGTGGTTGCGGAAATTGTGTTGTGTGCCCTGAACAGACTGAAGA |
| ONR1171 | TTGACAATTAATCATCGGCTCGTATAATGTTGGCCACCTCCAAACTAGA |

| | |
|---|---|
| ONR1162 | CTGATTGACGGCATCTTGCT |
| ONR1161 | AGACTAGGGAGCCAGCAATG |
| ONR1172 | GTCTTTGTCTTCTGCGTCGGGTTGATCGTGGTTGCGGAAATTGTGTTGTGTG ACGGTCGTTTTGTGTTGT |
| ONR1326 | CCTTTTCCATTTCTGAACATTGCCAATTAATGCAGCTGGCACGAC |
| ONR1281 | CGCCCATCAGGTAACGCGGTTTGTCCATTGTTCATTCCACGGACA |
| ONR1327 | AACCTGTCGTGCCAGCTGCATTAATTGGCAATGTTCAGAAATGGA |
| ONR1167 | GGTGGAATACGGTCTGGTCGATTCGCCACGCACGTTGTGATATG |
| ONR1168 | CTCACGAGAATCTTCCTGCGTACGCCATTGTTCATTCCACGGACA |
| ONR1169 | GTTTTTGTCCGTGGAATGAACAATGGCGTACGCAGGAAGATTCTC |
| ONR1170 | CATCTACATATCACAACGTGCGTGGCGAATCGACCAGACCGTATT |
| ONR1332 | GGTGGAATACGGTCTGGTCGATTCGATTAATGCAGCTGGCACGAC |
| ONR0001 | GTTTTCCCAGTCACGACGTT |
| ONR0002 | GCCTGATGCGGTATTTTCTC |
| ONR1057 | AACGTCGTGACTGGGAAAACCCACGCACGTTGTGATATGT |
| ONR1058 | GAGAAAATACCGCATCAGGCCCATTGTTCATTCCACGGAC |
| ONR1322 | CCACGCACGTTGTGATATGT |
| ONR1323 | CATAGCTGTTTCCTGTGTGA |
| ONR1324 | TCACACAGGAAACAGCTATGCAGTTTAAGGTTTACACCT |
| ONR1325 | ACATATCACAACGTGCGTGGTGGCTGTGTATAACGGAGCC |
| ONR0593 | TCTGACGCTCAGTGGAACGACCACGCACGTTGTGATATGT |
| ONR0594 | GCATTGGTAACTGTCAGACCCCATTGTTCATTCCACGGAC |
| ONR0597 | GCATTGGTAACTGTCAGACCCCACGCACGTTGTGATATGT |

| | |
|---|---|
| ONR0598 | TCTGACGCTCAGTGGAACGACCATTGTTCATTCCACGGAC |
| ONR1157 | AACGTCGTGACTGGGAAAACGAATGCTCCTGCTCATCTCC |
| ONR1158 | GAGAAAATACCGCATCAGGCTGACGGTCGTTTTGTGTTGT |
| ONR854 | CGAAACTGAAACAGGGCATT |
| ONR857 | TGCCCAACAGTAGCACTCAC |
| ONR1270 | TCAGGTCAATGCCGATAACA |
| ONR1271 | GATCTGCCGCTTTAGCATTC |
| ONR1205 | AATAGTGCTGGCACCCAAAC |
| ONR1206 | CCGCCTTCATTGATCTTTGT |
| ONR1195 | CGGTGATGTGACAAAACTGG |
| ONR1196 | TTATTGATGGGATCGGGTGT |
| ONR1316 | TATCTTTAACGGCGGTCTGG |
| ONR1317 | GCCTTCAACCAGGTCTTCTG |
| ONR1391 | AAGCGATTCGTCACTTTGCT |
| ONR1415 | GCCGTTATCGTCTGTTTGTG |
| ONR1416 | CATATCGGTGGTCATCATGC |

**Table S4: Primer sets and templates for DNA fragment preparation**

| **Name** | **size (bp)** | **Forward primer** | **Reverse primer** | **template** |
|---|---|---|---|---|
| cos(yajR) | 316 | ONR1280 | ONR1281 | λ cI857 genome |
| cat(vio) | 1091 | ONR1278 | ONR1099 | pKD3 |
| vioA | 1634 | ONR1102 | ONR1105 | *C. violaceum* JCM1249 genome |

| | | | | |
|---|---|---|---|---|
| vioAB | 4630 | ONR1102 | ONR1104 | *C. violaceum* JCM1249 genome |
| vio | 8237 | ONR1102 | ONR1103 | *C. violaceum* JCM1249 genome |
| yajR_L | 18684 | ONR1282 | ONR1097 | *E. coli* SN1171 genome |
| yajR_R | 18119 | ONR1100 | ONR1283 | *E. coli* SN1171 genome |
| cat(yajR) | 1116 | ONR1278 | ONR1163 | pKD3 |
| cat-vioA | 2675 | ONR1278 | ONR1105 | cat(vio) and vioA |
| cat-vioAB | 5671 | ONR1278 | ONR1104 | cat(vio) and vioAB |
| cat-vio | 9278 | ONR1278 | ONR1103 | cat(vio) and vio |
| cat(tac) | 1095 | ONR1278 | ONR1284 | pKD3 |
| vio(tac) | 8140 | ONR1272 | ONR1279 | *C. violaceum* JCM1249 genome |
| cos(yieL) | 346 | ONR1262 | ONR1263 | λ cI857 genome |
| cat(yieL) | 1085 | ONR1268 | ONR1284 | pKD3 |
| vio(yieL) | 8130 | ONR1166 | ONR1269 | *C. violaceum* JCM1249 genome |
| yieL_L | 15753 | ONR1264 | ONR1265 | *E. coli* SN1171 genome |
| yieL_R | 16629 | ONR1266 | ONR1267 | E. *coli* SN1171 genome |
| cos(ypjA) | 346 | ONR1201 | ONR1202 | λ cI857 genome |
| cat(ypjA) | 1085 | ONR1203 | ONR1284 | pKD3 |
| vio(ypjA) | 8130 | ONR1166 | ONR1204 | *C. violaceum* JCM1249 genome |
| ypjA_L | 15993 | ONR1197 | ONR1198 | *E. coli* SN1171 genome |
| ypjA_R | 15835 | ONR1199 | ONR1200 | *E. coli* SN1171 genome |
| cos(yneO) | 346 | ONR1191 | ONR1192 | λ cI857 genome |
| cat(yneO) | 1085 | ONR1193 | ONR1284 | pKD3 |
| vio(yneO) | 8130 | ONR1166 | ONR1194 | *C. violaceum* JCM1249 genome |

| | | | | |
|---|---|---|---|---|
| yneO_L | 16209 | ONR1187 | ONR1188 | *E. coli* SN1171 genome |
| yneO_R | 15686 | ONR1189 | ONR1190 | *E. coli* SN1171 genome |
| ccdBcos(yajR) | 819 | ONR1326 | ONR1281 | pUC19-ccdBcos |
| yajR(ccdB)_R | 18119 | ONR1100 | ONR1327 | *E. coli* SN1171 genome |
| lux | 6772 | ONR1173 | ONR1174 | *A. fischeri* JCM18803 genome |
| cos(yajR)2 | 316 | ONR1167 | ONR1168 | λ cI857 genome |
| yajR_L2 | 11316 | ONR1169 | ONR1097 | *E. coli* SN1171 genome |
| yajR_R2 | 11525 | ONR1100 | ONR1170 | *E. coli* SN1171 genome |
| pig_a | 11301 | ONR1171 | ONR1162 | *S. marcescens* JCM1239 genome |
| pig_b | 9924 | ONR1161 | ONR1172 | *S. marcescens* JCM1239 genome |
| ccdBcos(yajR)2 | 819 | ONR1332 | ONR1168 | pUC19-ccdBcos |
| yajR(ccdB)_R2 | 11525 | ONR1100 | ONR1333 | *E. coli* SN1171 genome |
| pUC19 | 2565 | ONR0001 | ONR0002 | pUC19 |
| cos | 307 | ONR1057 | ONR1058 | λ cI857 genome |
| pUC19-cos | 2724 | ONR1322 | ONR1323 | pUC19-cos |
| ccdB | 369 | ONR1324 | ONR1325 | JM109 genome |
| pUC19(cos) | 2573 | ONR0593 | ONR0594 | pUC19 |
| cos(pUC) | 307 | ONR0597 | ONR0598 | λ cI857 genome |
| pUCcos | 2719 | ONR0001 | ONR0002 | pUCcos |
| pig_a(pUCcos) | 11623 | ONR1157 | ONR1162 | *S. marcescens* JCM1239 genome |

| | | | | |
|---|---|---|---|---|
| pig_b(pUCcos) | 9894 | ONR1161 | ONR1158 | *S. marcescens* JCM1239 genome |

**Table S5: Primer sets for colony PCR**

| Purpose | Forward primer | Reverse primer |
|---|---|---|
| *yajR* check | ONR854 | ONR857 |
| *yieL* check | ONR1270 | ONR1271 |
| *ypjA* check | ONR1205 | ONR1206 |
| *yneO* check | ONR1195 | ONR1196 |
| *cos* check | ONR1316 | ONR1317 |
| *yajR-pig* check | ONR1391 | ONR857 |
| *ccdB* check | ONR1415 | ONR1416 |

**Table S6: Combinations of PCR fragments for PHAGE**

| Name | Fragment 1 | Fragment 2 | Fragment 3 | Fragment 4 | Fragment 5 | Fragment 6 |
|---|---|---|---|---|---|---|
| cat | cos(yajR) | yajR_L | cat(yajR) | yajR_R | | |
| cat-vioA | cos(yajR) | yajR_L | cat(vio) | vioA | yajR_R | |
| cat-vioAB | cos(yajR) | yajR_L | cat(vio) | vioAB | yajR_R | |
| cat-vio | cos(yajR) | yajR_L | cat(vio) | vio | yajR_R | |
| cat-tac-vio (yajR) | cos(yajR) | yajR_L | cat(tac) | vio(tac) | yajR_R | |

| | | | | | | |
|---|---|---|---|---|---|---|
| cat-tac-vio (yieL) | cos(yieL) | yieL_L | cat(yieL) | vio(yieL) | yieL_R | |
| cat-tac-vio (ypjA) | cos(ypjA) | ypjA_L | cat(ypjA) | vio(ypjA) | ypjA_R | |
| cat-tac-vio (yneO) | cos(yneO) | yneO_L | cat(yneO) | vio(yneO) | yneO_R | |
| cat-tac-vio_ccdB | ccdBcos(yajR) | yajR_L | cat(tac) | vio(tac) | yajR(ccdB)_R | |
| cat-tac-lux | cos(yajR) | yajR_L | cat(tac) | lux | yajR_R | |
| cat-tac-pig | cos(yajR)2 | yajR_L2 | cat(tac) | pig_a | pig_b | yajR_R2 |
| cat-tac-pig_ccdB | ccdBcos(yajR)2 | yajR_L2 | cat(tac) | pig_a | pig_b | yajR(ccdB)_R2 |

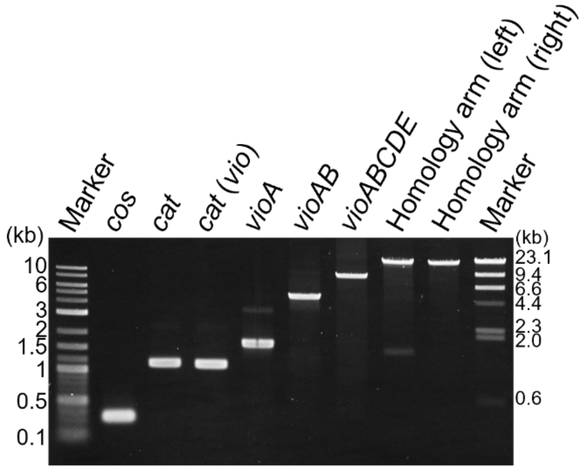


**Figure S1. Agarose gel electrophoresis of PCR fragments used in Fig. 2.**

PCR products corresponding to the constructs used in Fig. 2 were separated on a 1% agarose gel and visualized by ethidium bromide staining. Molecular weight markers were loaded on both sides of the gel, and the central lanes contain PCR products for *cos*, *cat*, *cat(vio)*, *vioA*, *vioAB*, *vioABCDE*, and the left and right homology arms, as indicated.

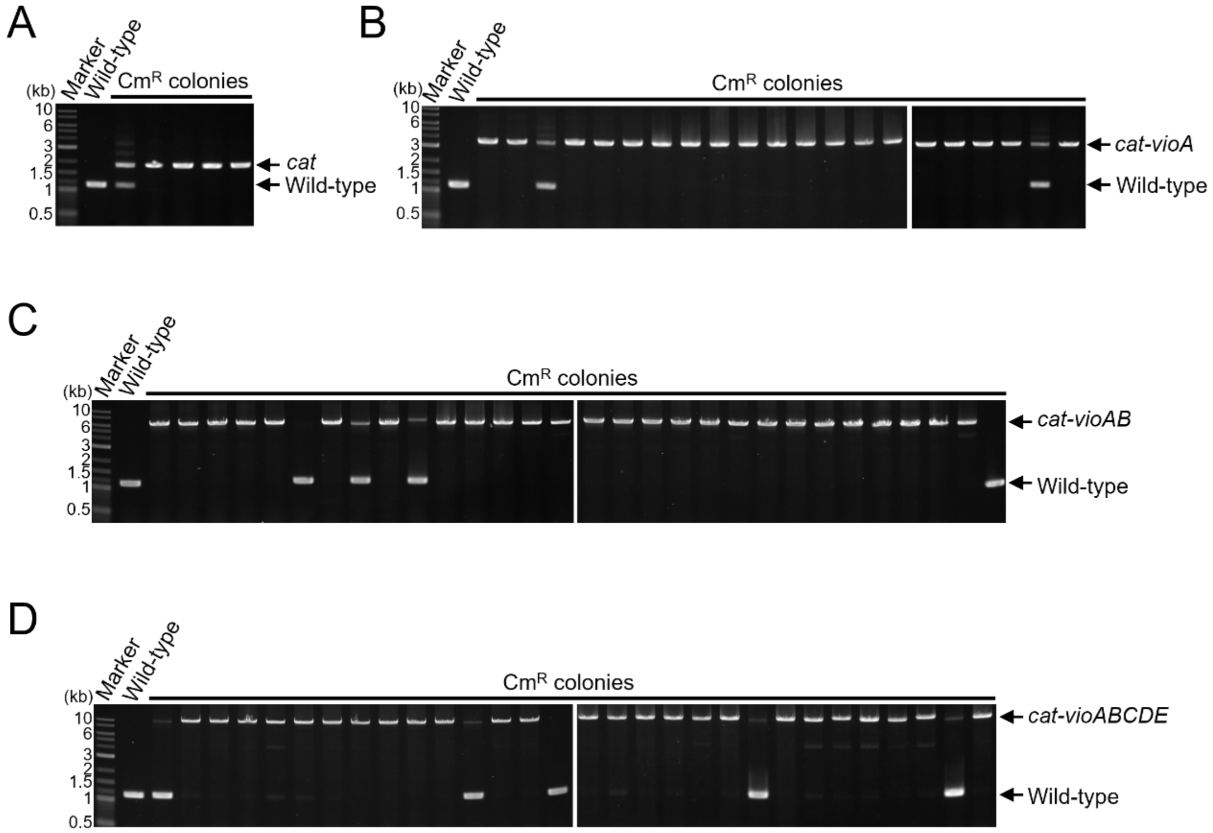


**Figure S2. Agarose gel electrophoresis of colony PCR products for constructs used in Fig. 2.**

Colony PCR was performed to verify the presence of the indicated integrations in chloramphenicol-resistant ($Cm^R$) colonies, and the resulting PCR products were analyzed by agarose gel electrophoresis. Panels A-D show PCR products corresponding to *cat* (A), *cat-vioA* (B), *cat-vioAB* (C), and *cat-vioABCDE* (D). In each panel, a wild-type strain was included as a negative control, and the adjacent lanes contain PCR products from individual $Cm^R$ colonies. The arrows in each panel indicate the bands corresponding to the expected insertion and wild-type allele.

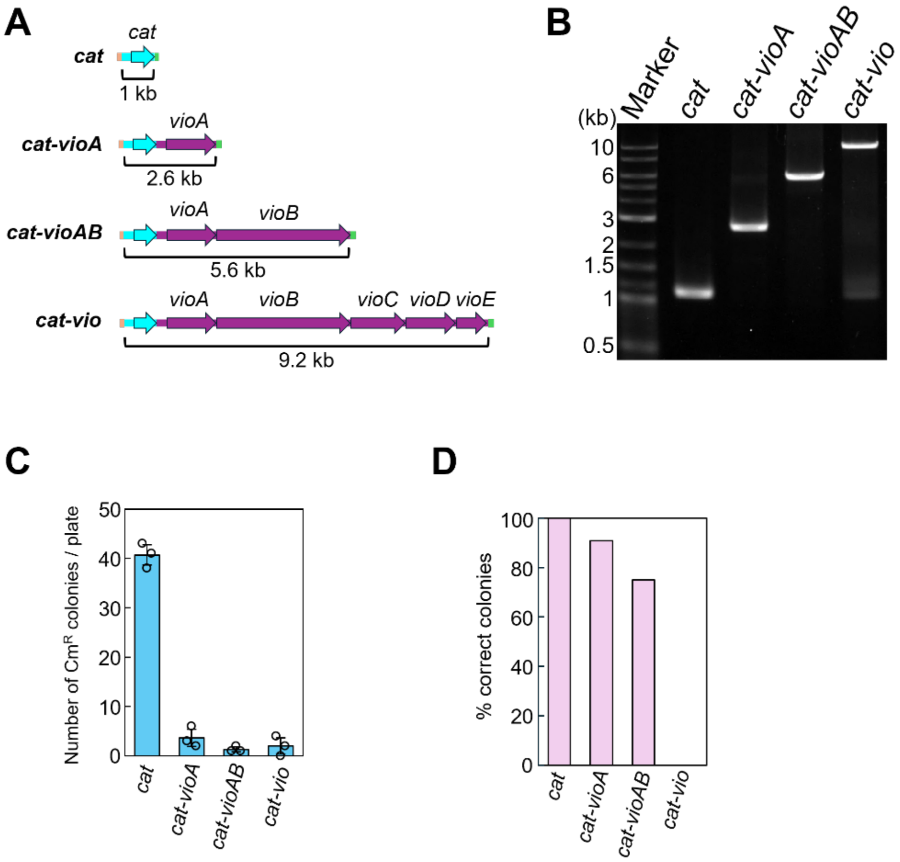


**Figure S3. Analysis of λ Red-mediated integration of *vio* gene constructs.**

(A) Schematic representation of the PCR fragments used for λ Red recombination. Four constructs (*cat*, *cat-vioA*, *cat-vioAB*, and *cat-vioABCDE*) were amplified with 50-bp homology arms appended to both ends to facilitate recombination into the target genomic locus.

(B) Agarose gel electrophoresis of the PCR fragments shown in panel A.

(C) Number of chloramphenicol-resistant ($Cm^R$) colonies obtained after λ Red-mediated recombination with each construct. Error bars represent standard deviations, and open circles indicate individual measurements.

(D) Proportion of $Cm^R$ colonies that correctly integrated the intended construct into the genome, as determined by colony PCR.

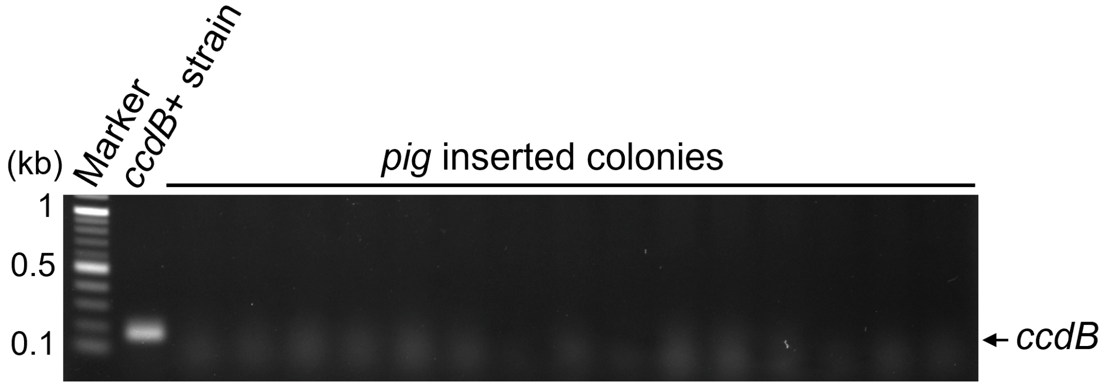


**Figure S4. Confirmation of the absence of *ccdB* by colony PCR**

Colony PCR was performed to determine whether the *ccdB* gene remained in the genome after recombination. The *ccdB*-containing strain XL10-Gold was included as a positive control. Fifteen colonies corresponding to the *ccdB*-based integration system shown in Fig. 7F were analyzed by colony PCR. The PCR products were separated by agarose gel electrophoresis, and the position corresponding to the PCR product derived from the *ccdB* target region is indicated by an arrow.

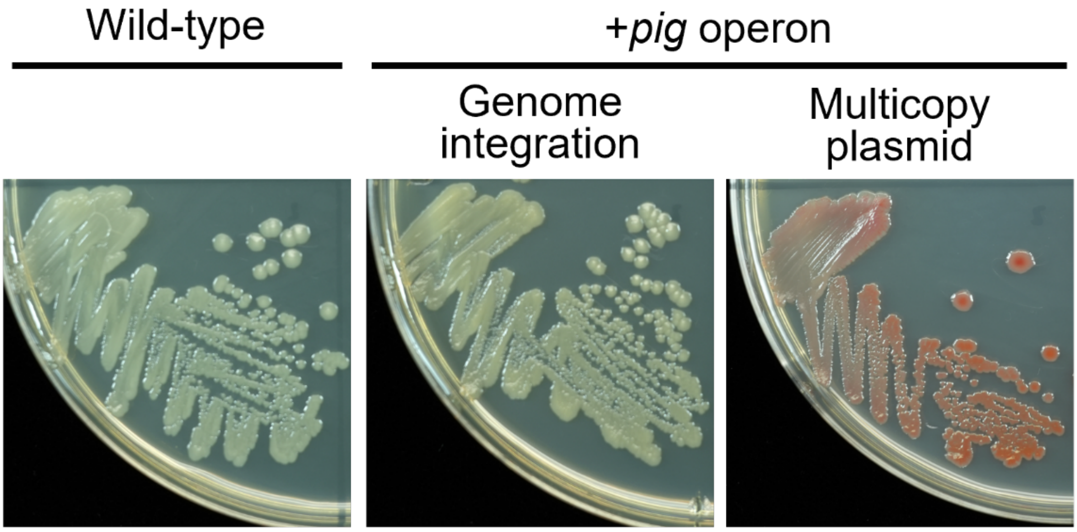


**Figure S5. Colony appearance of *E. coli* strains carrying the *pig* operon.**

Photographs of colonies grown on LB agar plates to assess pigment production resulting from the introduction of the *pig* operon. Plates were incubated at 25 °C for 3 days before imaging. A wild-type strain lacking the *pig* operon was included as a negative control. In addition to the wild-type strain, two types of strains carrying the *pig* operon are shown, as indicated: a strain in which the *pig* operon was integrated into the genome, and a strain carrying a multicopy plasmid pUCcos-pig harboring the *pig* operon.
